# Lateral gene transfer shapes the distribution of RuBisCO among Candidate Phyla Radiation bacteria and DPANN archaea

**DOI:** 10.1101/386292

**Authors:** Alexander L. Jaffe, Cindy J. Castelle, Christopher L. Dupont, Jillian F. Banfield

## Abstract

Ribulose-1,5-bisphosphate carboxylase/oxygenase (RuBisCO) is considered to be the most abundant enzyme on Earth. Despite this, its full diversity and distribution across the domains of life remain to be determined. Here, we leverage a large set of bacterial, archaeal, and viral genomes recovered from the environment to expand our understanding of existing RuBisCO diversity and the evolutionary processes responsible for its distribution. Specifically, we report a new type of RuBisCO present in Candidate Phyla Radiation (CPR) bacteria that is related to the archaeal Form III enzyme and contains the amino acid residues necessary for catalytic activity. Genome-level metabolic analyses supported the inference that these RuBisCO function in a nucleotide-based, CO_2_-incorporating pathway. Importantly, some Gottesmanbacteria (CPR) also encode a phosphoribulokinase that may augment carbon metabolism through a partial Calvin-Benson-Bassham Cycle. Based on the scattered distribution of RuBisCO and its discordant evolutionary history, we conclude that this enzyme has been extensively laterally transferred across the CPR bacteria and DPANN archaea. We also report RuBisCO-like proteins in phage genomes from diverse environments. These sequences cluster with proteins in the Beckwithbacteria (CPR), implicating phage as a possible mechanism of RuBisCO transfer. Finally, we synthesize our metabolic and evolutionary analyses to suggest that lateral gene transfer of RuBisCO may have facilitated major shifts in carbon metabolism in several important bacterial and archaeal lineages.

## INTRODUCTION

Forms I and II Ribulose-1,5-bisphosphate carboxylase/oxygenase (RuBisCO) are central to carbon fixation via the Calvin-Benson-Bassham (CBB) Cycle in algae, plants, and some bacteria. Forms III and II/III RuBisCO, discovered in Archaea, are believed to add CO_2_ to ribulose 1,5-bisphosphate (RuBP) involving a two-step reaction from nucleotides like adenosine monophosphate (AMP) [1]. These “bona-fide” RuBisCO enzymes were historically considered to be domain specific. In contrast, RuBisCO-like proteins (Form IV) in both Bacteria and Archaea perform functions distinct from carbon fixation and may be involved in methionine and sulfur metabolism [2]. Alongside enzymatic characterization, phylogenetic analyses suggested that the modern distribution of the RuBisCO superfamily could be explained by an archaeal origin and subsequent transfer by both vertical and lateral processes [3].

Recently, metagenomic studies of diverse environments introduced additional complexity to evolutionary considerations by uncovering new RuBisCO diversity. First, new genomes from Candidate Phyla Radiation (CPR) bacteria and DPANN archaea were shown to contain a hybrid II/III RuBisCO similar to that found in the archaeon *Methanococcoides burtonii* [4,5]. One version of this enzyme was shown to be catalytically active when expressed in a heterologous host [4,6]. An additional RuBisCO form related to the archaeal Form III, called the “III-like,” was reported from genomes of other members of the CPR and some DPANN archaea, further expanding enzyme diversity in these groups [5–7]. The III-like enzyme is also predicted to function in the CO_2_-incorporating AMP pathway. [6].

The discovery of RuBisCO in some CPR bacteria and DPANN archaea was interesting because these organisms have small genomes and lack many core metabolic functions. The common absence of pathways for synthesis of nucleotides, amino acids, and lipids has led to speculation that they live symbiotic or syntrophic lifestyles [4,8]. Despite this, recent research has shown that some members have a wide range of fermentative capabilities and the potential to play roles in carbon, nitrogen, sulfur, and hydrogen cycling [4,9]. A similar putative ecology has been suggested for the DPANN archaea [5]. The recovery of RuBisCO, functioning in an AMP pathway, expanded possible metabolic modes for some of these organisms and suggested that certain CPR and DPANN could derive energy and/or resources from ribose produced by other community members [6,7].

Major outstanding questions concern the full scale of RuBisCO diversity and how the various forms are distributed phylogenetically within the CPR and DPANN groups. Importantly, in the last few years, additional work has recovered CPR and DPANN genomes from a much wider array of environmental types, including additional groundwater locations, the deep subsurface, hydrocarbon-impacted environments, and the ocean [10–12]. Here, we examined over 300 genomes from metagenomes from these environments to further elucidate diversity, potential functions, and the evolutionary history of RuBisCO in members of the bacterial Candidate Phyla Radiation and DPANN archaea. First, we expand the distribution of forms to new phylogenetic groups and propose a major new type that, in some cases, might act in concert with phosphoribulokinase to augment carbon metabolism. Additionally, we describe a clade of putative RuBisCO-like proteins encoded by bacteriophage from diverse environments. Drawing on these observations and previous analyses, we suggest that lateral gene transfer may have been largely responsible for patterns of distribution of this enzyme. These lateral transfers could, in the presence of genes from other pathways, introduce new RuBisCO-based metabolic capacity to CPR and DPANN lineages.

## RESULTS

### Metagenomics expands the diversity of RuBisCO forms and reveals putative new enzyme types in CPR and phage

The large majority of CPR and DPANN RuBisCO sequences analyzed fell into five clearly defined phylogenetic groups **(Fig. 1)**, four of which (II/III, III, III-like, IV) correspond to the “Forms” defined in previous literature:

**Figure 1.**
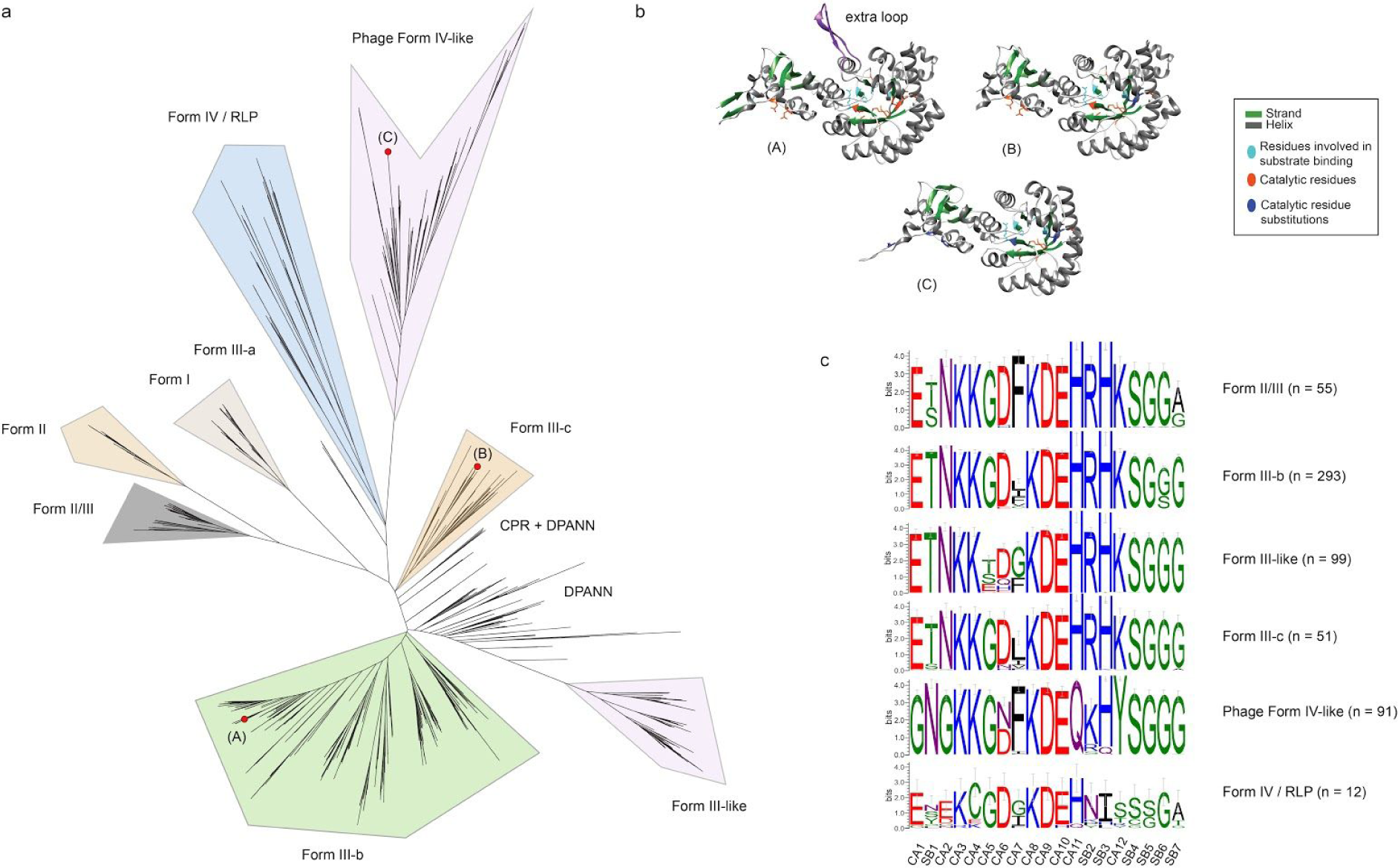
Known RuBisCO diversity is expanded by metagenomics. **(a)** Maximum-likelihood tree for de-replicated RuBisCO large-chain sequences, delineated by “Form.” (A), (B), (C) show the phylogenetic position of protein sequences modeled in **(b). (c)** Aggregate sequence logos for protein sequences in each phylogenetic “Form,” describing key catalytic activity (CA) and substrate binding (SB) residues in RuBisCO sequences. Logo colors represent residues with similar chemical properties.

#### II/III

Our analysis recovered new sequences of the Form II/III RuBisCO, originally thought to be exclusively found in Archaea [13]. Here, we broaden the phylum-level distribution of the group with additional representatives from the DPANN groups Woesearchaeaota and Micrarchaeota **(Fig. 2)**. Form II/III RuBisCO of CPR and DPANN partition into two subgroups based on the presence or absence of a 29 amino acid (or longer) insertion, the biochemical implications of which are currently unknown [6]. All but one of the full-length Micrarchaeaota and Woesearchaeota sequences recovered in this study contained insertions in the expected region.

**Figure 2.**
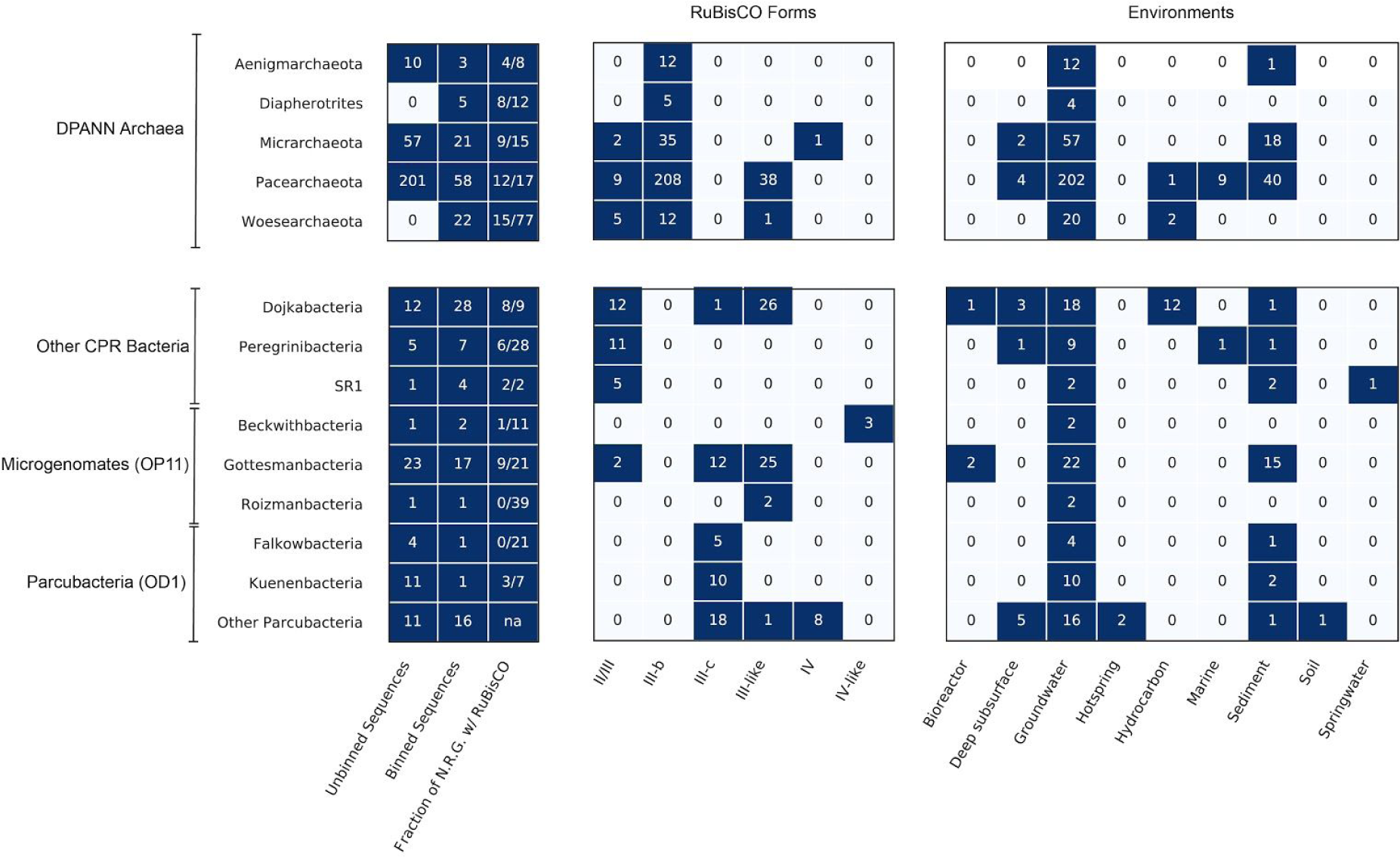
RuBisCO diversity among the CPR bacteria and DPANN archaea. Boxes represent counts of de-replicated protein sequences used in the phylogenetic analysis (Unbinned and Binned Sequences) as well as the number of those sequences that fell into described RuBisCO Forms and Environments. Fraction of non-redundant genomes (N.R.G.) with RuBisCO describes the proportion of genomes per phylum encoding RuBisCO after dereplication at 99% ANI (see Methods).

#### III

We also identified archaeal Form III RuBisCO sequences in a variety of DPANN, and, potentially, CPR genomes. We refer to this as Form III-b to distinguish it from Form III-a, a divergent group described by Kono et al. [14] and the Form III-like proteins [6]. Most of the newly reported Form III-b sequences contained the critical substrate binding and catalytic residues for RuBisCO function but also largely shared residue identity with canonical archaeal versions **(Fig. 1)**. Notably, we found this enzyme form in three DPANN groups - the Diapherotrites, the Micrarchaeota, and the Woesearchaeota - previously not known to harbor Form III-b RuBisCO, extending the presence of these enzymes beyond Pacearchaeota and Aenigmarchaeota [5], two major groups in the DPANN **(Fig. 2)**. Approximately 70 DPANN sequences were outside the clade containing characterized Form III-b, but were assigned to this type given low support for separating branches and overall closer relatedness to III-b than III-like sequences (DPANN (Form III-b), **Fig.2)**.

We identified several previously published sequences assigned to genomes from the Levybacteria and Amesbacteria phyla (Microgenomates superphylum) deeply nested within the archaeal Form III-b sequences. In addition, we recovered a set of unbinned sequences with highest similarity to these same binned Levybacteria and Amesbacteria genomes. The Levybacteria sequences group loosely with sequences from Aenigmarchaeota and a reference sequence from the archaeon *Methanoperedens nitroreducens* (∼80% identity). In contrast, the Amesbacteria sequences grouped most closely to those from a clade from Pacearchaeota. To verify the binning of these genome fragments, we examined the top BLAST hits to well annotated genes on each genome fragment bearing a RuBisCO gene. For the fragment putatively assigned to Amesbacteria, BLAST affiliation of non-hypothetical genes was inconclusive - some genes had only low identity with archaeal genes, others had identity to both bacterial and archaeal genes, while one gene encoding a serine acetyltransferase had ∼ 70% bacterial identity. In addition to RuBisCO, the ∼34 kbp putative Levybacteria genome fragment encodes a large CRISPR-Cas locus, a novel transposase and a phage/plasmid primase, all of which may indicate mobile genetic material. BLAST affiliation of well-annotated genes was also inconclusive for this fragment - if not of bacterial origin, the genome fragment could be phage or plasmid. Future research is needed to confirm that Form III-b RuBisCO is encoded in genomes of some CPR bacteria.

#### III-like

Many RuBisCO sequences placed phylogenetically in a deeply-branching, monophyletic clade that is divergent from the archaeal Form III and referred to as “III-like” [6] **(Fig. 1)**. We added 12 Dojkabacteria (WS6, a group within the CPR) sequences from hydrocarbon-impacted environments and nearly 40 DPANN sequences from Woesearchaeota and Pacearchaeota from multiple environments **(Fig. 2)**. As reported previously [6], the “III-like” sequences recovered in this analysis appear to have insertions mostly 11 amino acids in length.

#### III-c

We report a deeply-branching clade of RuBisCO sequences not described in previous work **(Fig. 1)**. While overall sequence similarity is closest to described Form III-b RuBisCO (∼50%), our analysis clearly indicates a distinct phylogenetic status. This group, which we term “Form III-c”, contained ∼50 sequences from the Dojkabacteria and several groups in the Microgenomates and Parcubacteria superphyla. Form III-c was particularly abundant in the Gottesmanbacteria and Kuenenbacteria, the latter of which appears to harbor this form exclusively **(Fig. 2)**. We identified organisms bearing this enzyme type in almost every environment included in our study, indicating that it may compose an important and previously unrecognized aspect of RuBisCO diversity. To predict the biochemical potential of this type, we examined 12 residues known to be important for catalytic activity and 7 important for substrate binding from each sequence [2,15]. This analysis indicated that Form III-c RuBisCO sequences contain critical residues largely identical to those reported for Form III-b enzymes; however, a minority of sequences encoded modifications to catalytic site #6 that could alter its chemical properties **(Fig. 1c)**. Additionally, modeling through I-TASSER revealed that secondary structure of an III-c sequence was consistent with that of existing Form III-b templates **(Fig. 1b)**.

#### IV and IV-like

Form IV (RLP) proteins are a clade of highly divergent RuBisCO with low sequence similarity (∼30%) to ‘bona fide’ RuBisCO enzymes and divergent functions [2,16]. Our phylogenetic analyses identified 8 Form IV (RLP) RuBisCOs in CPR genomes, all in the genomes of bacteria from the Parcubacteria superphylum **(Fig. 2)**. The Parcubacteria Form IV sequences compose a monophyletic clade nesting within the previously described IV-Photo type RLP. However, the key catalytic/substrate binding residues are only partially conserved **(Fig. 1c)**. We only recovered one RLP from a Micrarchaeota (DPANN) genome **(Fig. 2)**, despite the fact that archaeal sequences appear to be at the root the RuBisCO/RLP superfamily [3].

Using manually curated HMMs, we also identified ∼90 non-redundant sequences related to Form IV RuBisCO on putative phage as well one unusually large, curated phage genome (∼ 200 kbp) **(Fig. S1a)**. The phage sequences appeared highly divergent from other RLPs, but most shared some key residues found in canonical Form IV RuBisCOs **(Fig. 1c)** and scored highly on HMMs constructed from verified Form IV RuBisCO (score > 200, e-val << 0.05). Additionally, modeling of one sequence revealed a secondary structure consistent with large chain templates but missing several conserved residues, as expected for Form IV enzymes **(Fig. 1b)**. Analysis of co-encoded phage terminase proteins indicated that at least some of the viral sequences were from members of the Myoviridae. Three other RLP-related sequences attributed to Beckwithbacteria grouped with those from putative/confirmed phage and together appeared as an outgroup to all other RLPs **(Fig. 1a)**. Analysis of the genes encoded on the manually curated Beckwithbacteria fragment with IV-like RuBisCO supported its bacterial origin **(Fig. S1b)**.

### Form III-c RuBisCO is encoded in close proximity to genes of the CO_2_-incorporating AMP pathway

To test the hypothesis that the Form III-c RuBisCO participates in the previously described AMP pathway, we narrowed our focus on the high-quality genomes containing this form of the enzyme. The AMP pathway employs AMP phosphorylase (*deoA*) and R15P isomerase (*e2b2*) to provide RuBisCO with RuBP substrate, incorporating CO_2_ to produce two 3-Phosphoglyceric acid molecules (3-PGA) which are fed into upper glycolysis. Forty-one genomes encoding Form III-c RuBisCO (68%) also contained homologs for *e2b2* and *deoA*. Most Gottesmanbacteria (OP11) contained homologs to *deoA* at a threshold lower than originally used in the KEGG analysis, as these sequences generally had a conserved deletion of ∼80 amino acids at the start of the protein compared to other CPR proteins. Despite this, we predict that these proteins are divergent *deoA* homologs based on phylogenetic placement and re-alignment with reference sequences. Among Gottesmanbacteria genomes, the percentage containing all three genes associated with AMP metabolism was higher (∼92%). Thus, the new Form III-c RuBisCO, especially among Gottesmanbacteria, is consistently associated with *deoA* and *e2b2* homologs, as reported previously for CPR genomes with other forms of the enzyme [6] **(Fig. S2a)**.

Next, we analyzed the genomic proximity of *deoA* and *e2b2* homologs with RuBisCO in all genomes containing Forms III-b, III-c and II/III RuBisCOs. Forty-six genomes contained fragments encoding all three genes on the same stretch of assembled DNA **(Fig. S2b)**. Most fragments encoded RuBisCO between the other genes, although a minority of genomes appeared to contain rearrangements **(Fig. S2c)**, as previously reported for a Form II/III-bearing PER-1 genome [6]. Next, for cases where all three genes occurred on the same contig, we defined a metric called “pathway proximity” (sum of the genomic distance between the three genes, see Methods). This analysis suggested that genes involved in AMP metabolism are more frequently and more proximally co-located in the genomes of organisms bearing the Form III-c RuBisCO than in genomes with other forms, even though these RuBisCO were present on assembled fragments of similar length **(Fig. 3a)**. Two outlier Form III-c genomes from a previous study [12] encoded extremely distant *deoA* genes on long fragments, possibly the result of genetic rearrangement or errors in genome assembly. Interestingly, some genomes bearing Form III-c RuBisCO encoded the three genes consecutively. In other cases the RuBisCO gene was fused to the isomerase **(Fig. 3b, Fig. S3)**. The close proximity and fusion support the conclusion that this RuBisCO form participates in the CO_2_-incorporating AMP pathway.

**Figure 3.**
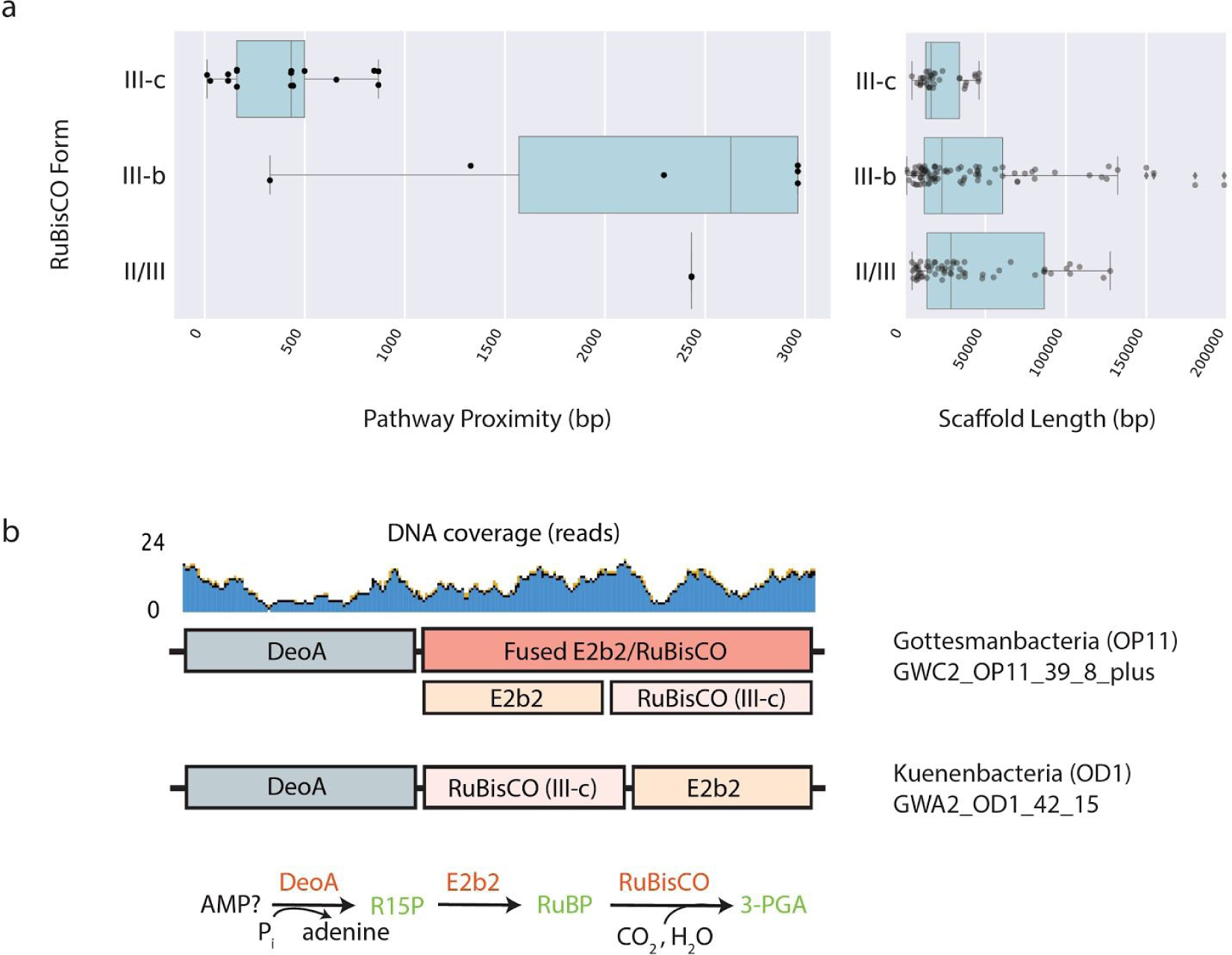
Genomic context of Form III-c RuBisCO in CPR and DPANN. **(a)** Proximity of genes on genome fragments encoding all three components of the CO_2_-incorporating AMP pathway, and the length of all
binned fragments containing the specified RuBisCO Form. **(b)** Genomic diagram of AMP components in CPR genomes with fused RuBisCO and consecutive ordering of genes on the chromosome.

### RuBisCO phylogeny is incongruent with that of e2b2 *and* deoA

The discovery of the Form III-c suggested that lateral gene transfer may play a role in shaping the distribution of this enzyme in CPR and DPANN phyla. To test for preliminary evidence of lateral gene transfer involving the genes of the AMP pathway, we compared the phylogeny of RuBisCO with that of the other pathway components [6]. We found that the phylogenies for CPR and DPANN *deoA* and *e2b2* recapitulate the organism phylogeny, yet were largely incongruent with that of RuBisCO **(Fig. S4)**. Among the monophyletic Gottesmanbacteria, genomes with the same RuBisCO Form (III-c, III-like) appeared to cluster together, as would be expected if lateral gene transfer of various RuBisCO followed vertical inheritance of *deoA* and *e2b2* by different sub-phylum level lineages. Although still undersampled, there is some indication that RuBisCO-correlated phylogenetic clustering will emerge for Parcubacteria (OD1) and Dojkabacteria (WS6), which at the phylum level contain high RuBisCO diversity.

### CPR bacteria genomes bearing Form III-c RuBisCO also encode phosphoribulokinase

We examined whether recovered CPR RuBisCO, including the Form III-c, might participate in a form of the CBB pathway by searching all binned genomes for homologs of phosphoribulokinase (PRK). PRK is a key CBB marker gene that is critical for regenerating RuBP substrate. Our analysis recovered 31 genomes encoding PRK homologs among the Gottesmanbacteria and Peregrinibacteria, the first PRK homologs reported in the CPR. PRK sequences were encoded by Gottesmanbacteria harboring Form III-like and III-c RuBisCO, whereas Peregrinibacteria PRK were associated with the typical Form II/III enzyme. Phylogenetic analysis revealed that Gottesmanbacteria PRK sequences comprised a highly-supported monophyletic clade nested within “putative bacterial type” sequences, in particular several from recently isolated cyanobacterial genomes **(Fig. 4, Fig. S5)**. These putative cyanobacterial sequences appear to be distinct from classical cyanobacterial PRK and have not yet been assayed for functional activity. Similarly, two recovered Peregrinibacteria PRK form a monophyletic clade sister to additional divergent cyanobacterial sequences **(Fig. 4, Fig. S5)**. To test whether CPR genomes containing PRK have the full genomic repertoire for the CBB cycle, we searched genomic bins for nine other genes involved in this pathway (**Table S2**). Several Gottesmanbacteria bins contained near-complete CBB pathways, lacking only the gene for sedoheptulose 1,7 bisphosphatase (SBP, **08 in Fig. 4**). Additionally, instead of separate genes for fructose 1,6-bisphosphate aldolase (FBA) and fructose 1,6-bisphosphatase (FBPase), most Gottesmanbacteria genomes encoded an enzyme most similar to a bifunctional version found in some thermophilic, chemoautotrophic bacteria and archaea [17] **(Fig. 4)**. One Peregrinibacteria genome contained all genes involved in the CBB cycle with the exception of FBPase (06), although additional sampling and reconstruction of high quality genomes is necessary to definitively designate this enzyme as missing.

**Figure 4.**
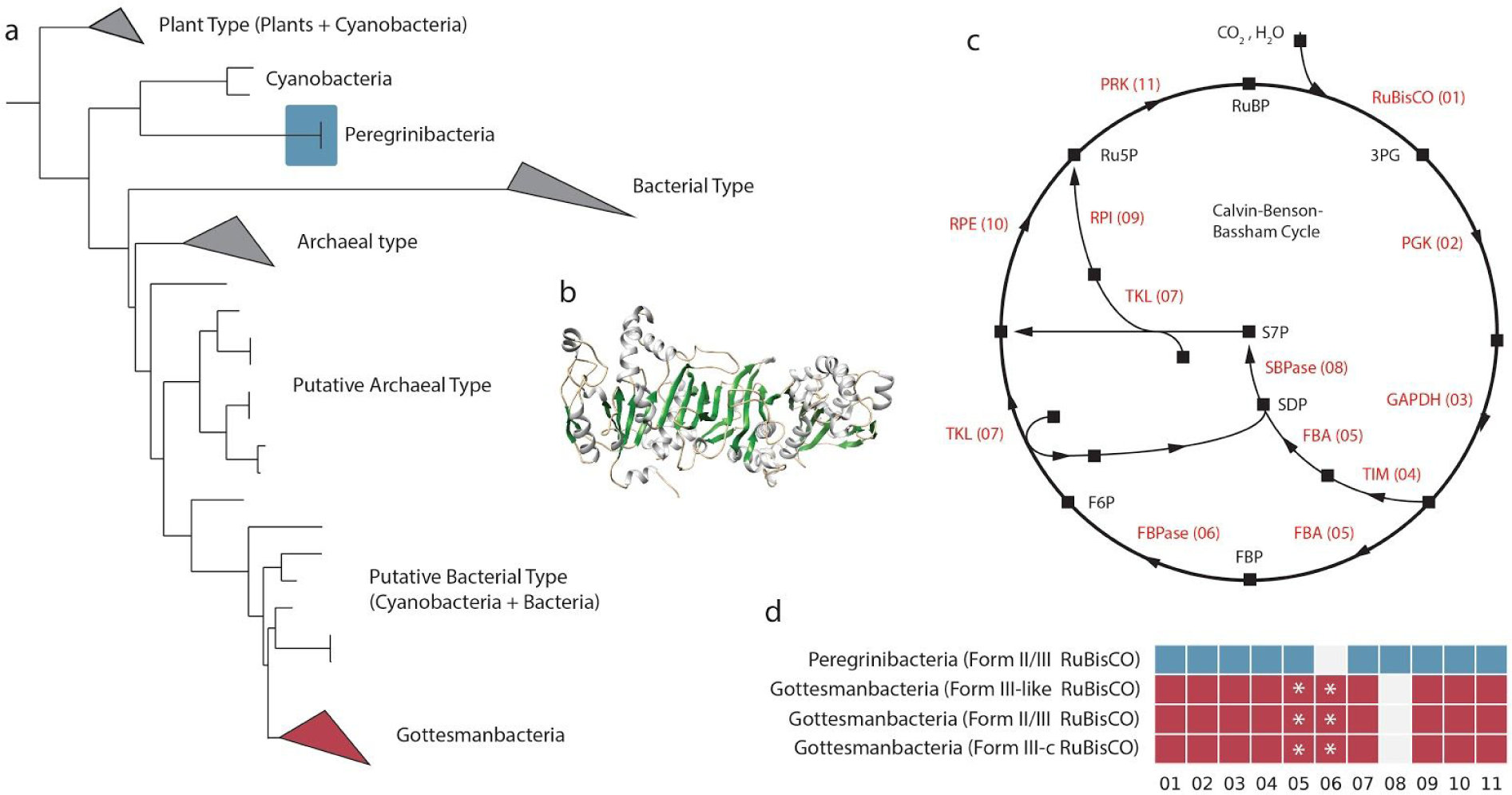
Some CPR bacteria encode a putative phosphoribulokinase (PRK). **(a)** Maximum-likelihood tree showing phylogenetic position of putative PRK in CPR phyla. See **Fig. S5** for fully labeled tree. **(b)** Example protein model of putative PRK in the CPR phylum Gottesmanbacteria. **(c)** Schematic of the Calvin-Benson-Bassham Cycle. Squares represent molecular intermediates, while arrows represent enzymatic steps labelled in red. Abbreviations: PGK, phosphoglycerate kinase; GAPDH, glyceraldehyde 3-phosphate dehydrogenase; TIM, triosephosphate isomerase; FBA, fructose-bisphosphate aldolase; FPBase, fructose 1,6-bisphosphatase; TKL, transketolase; SBPase, sedoheptulose-bisphosphatase; RPI, ribose-5-phosphate isomerase; RPE, ribulose 5-phosphate 3-epimerase. **(d)** Genomic repertoires of CPR bacteria encoding PRK. Numbers refer to enzymatic steps in (c). Asterisks indicate genomes that harbor an enzyme with highest homology to a bifunctional FBA/FBPase instead of separate FBA and FBPase.

## DISCUSSION

### New diversity in the RuBisCO superfamily and its implications for phylogenetic distribution and metabolic function among the CPR and DPANN

Through increased metagenomic sampling of diverse environments, we provide new information about the distribution of RuBisCO in major bacterial and archaeal groups and expand RuBisCO superfamily diversity. Our results also allow a quantitative assessment of RuBisCO diversity, revealing that the Pacearchaeota and Dojkabacteria in particular frequently encode various forms of the enzyme across many environmental types **(Fig. 2)**. This suggests that RuBisCO may be a core metabolic enzyme for these groups, which appear to have the most minimal metabolic and biosynthetic capacities among the DPANN and CPR radiations [7].

At present, Form III-c sequences occur only in several CPR lineages, contrasting with other Form III-related enzymes with archaeal representatives. Specifically, Form III-a is only known in methanogens, Form III-b is known to occur in archaea and possibly two CPR lineages, as well as another bacterium (*Ammonifex degensii*) [18], and Form III-like enzymes appear to be widely (but sparsely) distributed in both DPANN archaea and CPR bacteria. However, like the other Form III-related enzymes, the association of Form III-c RuBisCO with *e2b2* and *deoA* suggests this enzyme may also function in an AMP metabolism pathway. This pathway, originally described for the Form III-b enzyme in *Thermococcus kodakarensis*, relies on two proteins to provide RuBisCO with its substrate molecule, ribulose-1,5-bisphosphate (RuBP) [1]. First, an AMP phosphorylase encoded by the *deoA* gene catalyzes the release of ribose-1,5-bisphosphate (R15P) which is then subsequently converted to RuBP by a R15P isomerase encoded by *e2b2* [1]. Next, the RuBisCO incorporates H_2_O and CO_2_ with this substrate to create two molecules of 3-phosphoglycerate (3-PGA), which in turn can be diverted into central carbon metabolism [1]. Among the CPR, this pathway is thought to provide a simple mechanism for ribose salvage that may facilitate the syntrophic ecology of these organisms [6,7]. Contig-level analyses revealed a notable spatial association and occasional fusion of genes involved in AMP metabolism in genomes bearing the Form III-c RuBisCO, supporting the association of RuBisCO with this pathway. However, it is critical that future studies characterize the specific biochemistry of this new form and its possible function in CPR bacteria.

Form I and Form II RuBisCO function in the Calvin-Benson-Bassham cycle, which relies on phosphoribulokinase (PRK) to regenerate RuBP before carbon fixation by RuBisCO **(Fig. 4)**. The presence of PRK in Gottesmanbacteria and Peregrinibacteria raises the possibility that a CBB-like pathway may operate in carbon assimilation in these organisms. Lacking from their genomic repertoires, however, is sedoheptulose 1-7 bisphosphatase (SBPase) **(Fig. 4)**. In plants, SBPase catalyzes the phosphorylation of sedoheptulose 1,7 bisphosphate and is important for regulation of intermediate molecules in the CBB cycle [19]. Among Cyanobacteria, it has been shown that a single enzyme often functions as both an SBPase and a fructose bisphosphatase (FBPase), catalyzing a similar reaction on fructose bisphosphate in the second branch of the cycle **(Fig. 4)** [20,21]. Similarly, biofunctional activity has been demonstrated for other bacterial FBPase enzymes in both the CBB (*Ralstonia eutropha*) and a ribulose monophosphate cycle (*Bacillus methanolicus*) [22,23]. Complicating the possibility of a bifunctional FBPase/SBPase in the Gottesmanbacteria is the observation that most of these genomes encode a single enzyme most similar to a bifunctional fructose 1,6-bisphosphate aldolase (FBAase)/phosphatase, instead of separate FBPase and FBA. In the archaeal and bacterial lineages in which bifunctional FPA/FBPases have been characterized, these enzymes are thought to play a role in gluconeogenesis instead of the CBB pathway [17]. Thus, the association of this gene in the classical CBB in Gottesmanbacteria would require tripartite function as a FBPase, FBAase, and SBPase. As such, the functioning of a CBB-like pathway in CPR remains uncertain.

An alternative inference is that PRK contributes to carbon metabolism in Gottesmanbacteria by re-generating RuBP as substrate for RuBisCO in the AMP pathway. The same may be true of the two Peregrinibacteria genomes encoding the PRK but missing FBPase **(Fig. 4)**. In this scenario, components of the oxidative pentose phosphate pathway could convert glucose-6P into ribulose-5-P, which could then be converted to RuBP by the PRK **(Fig. S6)**. Going forward, it is critical that the PRK from both lineages, as well as the Gottesmanbacteria FBPase/FBA, be characterized biochemically, especially given that these sequences are divergent from well characterized enzymes **(Fig. S7)**. In any case, the presence of PRK, RuBisCO, and a putative bifunctional FBPase/FBA in CPR genomes suggests that these organisms may have acquired fundamental components of carbon metabolism by lateral transfer. This extends the prior observations of specific transfer of the bifunctional FBA/FBPase to bacteria [17] as well as the more general occurrence of transfer among CPR bacteria [24].

Finally, we report Form IV RuBisCO-like proteins (RLPs) in the genomes of several CPR bacteria and a DPANN archaeon. Previously described RLPs fall into 6 clades and have distinct patterns of active site substitutions that likely affect their functionality [3]. Sequences from Parcubacteria and Micrarchaeota were mostly closely related to a known RLP clade called the IV-Photo, which is implicated in sulfur metabolism/stress response in green sulfur bacteria like *C. tepidum [16]*. While active site residue divergence complicates the inference of function based on phylogenetic placement, it is possible that Form IV sequences found in CPR bacteria also function in oxidative stress response. Our analysis also revealed the presence of a large, highly divergent clade of RuBisCO sequences related to Form IV/RLP in the genomes of bacteriophage **(Fig. 1a)**, some of which were classified as Myoviridae. That these sequences were recovered from various environments suggests that these phage proteins may be a widespread and to date underappreciated reservoir of diversity in the RuBisCO superfamily. Sequence analysis revealed that these putative RuBisCO-like proteins encoded key residues highly divergent from known type IVs, leaving possible functionality unclear. However, future work may uncover “bona fide” RuBisCO-like proteins on phage genomes, supporting the inference that these enzymes are widely laterally transferred across lineages. Previous work has shown that marine phage can impact host carbon metabolism through auxilary expression of other photosynthetic genes [25,26]. Regardless of type, phage-associated proteins with homology to RuBisCO should be assessed functionally to evaluate whether they have to the potential to augment host metabolism in the natural environment.

### Sparse distribution of the RuBisCO superfamily suggests that lateral gene transfer shapes the distribution of multiple Forms among DPANN, CPR, and bacteriophage

Ancient lateral gene transfer has been invoked in several previous studies to explain the current distribution of RuBisCO in bacteria and archaea. This process probably drove the evolution of Form IV as well as Forms I and II from the ancestral Form III [2,3], ultimately resulting in diverse RuBisCO types that now occur in Archaea, Bacteria, and Eukaryotes. Results of the current study support this inference and extend it, suggesting that lateral gene transfer has also played a role in distributing recently recognized RuBisCO forms across Archaea and Bacteria. First, the discovery of Form III-c RuBisCO in CPR bacteria suggests gene transfer between CPR bacteria and archaea, as this new Form is most closely related to the archaeal Form III-b. The recovery of canonical archaeal Form III-b proteins within previously published Amesbacteria and Levybacteria bins (both CPR), if verified, would support this conclusion. Additionally, previous findings of a Dojkabacteria (WS6) genome harboring both Form III-like and Form II/III RuBisCO and a divergent Form III-b RuBisCO enzyme in the Firmicute *Ammonifex degensii* are best explained by gene acquisition via lateral transfer [10,18].

Broader evolutionary patterning supports the idea that lateral gene transfer has played an underappreciated role in the shaping the evolution of RuBisCO among CPR and DPANN. Our results reveal a relatively wide but sparse distribution of RuBisCO across CPR/DPANN lineages, with most lineages containing very low frequencies of the enzyme **(Fig. 2)**. The non-congruency of RuBisCO phylogeny with those of *deoA* and *e2b2*, which appear to have been largely vertically transmitted in CPR lineages **(Fig. S4),** suggests divergent evolutionary histories of these functionally related genes. Thus, we conclude that the distribution of RuBisCO diversity in both CPR and DPANN is more likely to be explained by LGT than extensive gene loss.

The results presented in this study give new insights on the evolution of the RuBisCO superfamily as a whole. **Fig. 5** is a schematic diagram that integrates ideas of Tabita and colleagues [3] with inferences arising from our results. Previous work has suggested that an ancestor of Methanomicrobia laterally transferred a Form III enzyme to a bacterial ancestor (Step 1 in Fig. 5) where it subsequently evolved to generate both the Form I and Form II enzymes (Step 2). The findings of the current and prior studies [3,6] indicate that an ancestral Form III sequence may have diverged from there into at least three Form III-related types (Step 3 and 4 in **Fig. 5),** including the Form III-b (traditional archaeal form) and the newly reported Form III-c. Specifically, we speculate that an additional transfer of the III-b enzyme from Archaea to a CPR led to the evolution of the III-like and III-c Forms (Steps 3, 4). Subsequent transfers to both the Dojkabacteria (Step 5), Parcubacteria (Step 6), and the common ancestor of Pacearchaeota and Woesearchaeota (Step 7) would recapitulate the current distribution of of III-like enzymes across the tree of life. However, with the current evidence, we cannot rule out the possibility that the III-like RuBisCO evolved within DPANN archaea and was transferred in the reverse direction to the CPR. Interestingly, Form III-like enzymes occur only in some CPR lineages and DPANN archaea, without any known representatives outside these radiations. Many DPANN lineages also encode the classical archaeal Form III-b, currently thought to have originated in the Methanomicrobia [3]. Recent phylogenomic studies of the Archaea have inferred a root in between DPANN and all other groups [27], requiring a transfer of Form III-b to DPANN from another lineage to explain this form’s extant distribution (Step 8).

**Fig. 5.**
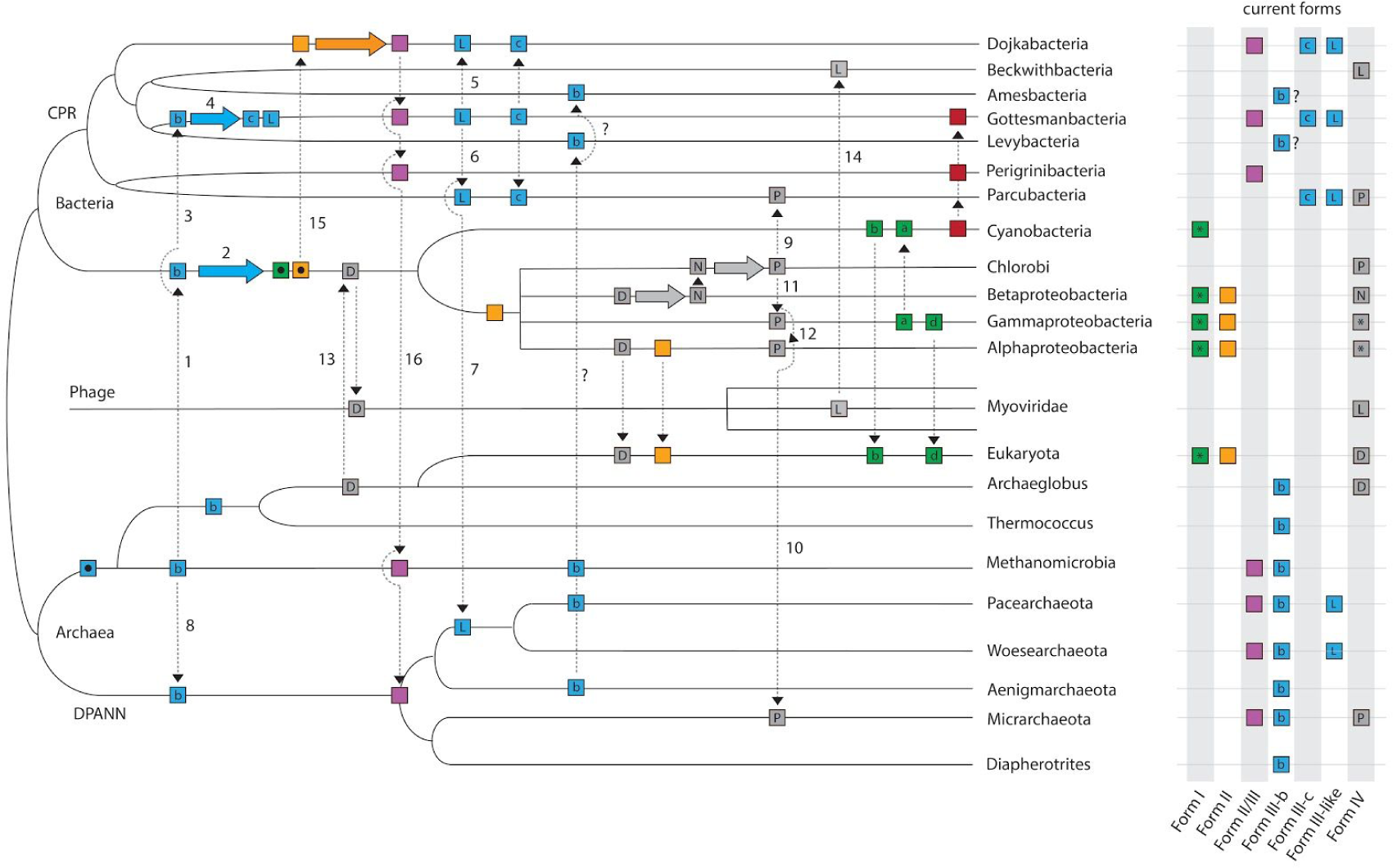
Conceptual diagram illustrating the role of lateral gene transfer in the evolution of RuBisCO and PRK (in red). Distinct RuBisCO Forms are represented by boxes of different colors, and Form subtypes are indicated by a letter within the box if applicable. Form III enzymes are expanded for additional clarity and are abbreviated as follows: b, Form III-b; c, Form III-c, L, Form III-like. Form IV abbreviations are as follows: D, DeepYkr; N, NonPhoto, P, Photo (see Tabita et al., 2007); L, Form IV-like (this study). Ancestral sequences are represented by black dots inside boxes. Dotted arrows represent possible lateral gene transfers, solid colored arrows represent evolution of RuBisCO within a lineage. Step numbers are referenced in the Discussion section. **N.B.** This tree does not convey time-calibrated information and is arranged to optimize conceptual understanding over accurate evolutionary relationships.

Our results also suggest the importance of lateral transfer processes in shaping the distribution of Form IV RuBisCO. For example, the Parcubacteria and Micrarchaeota Form IV enzymes are similar to those in characterized green sulfur bacteria, and may indicate transfer from this source (Steps 9,10 in **Fig. 5**). Previous work has suggested that the IV enzyme is mobile and may have been transferred from Chlorobi to both Gammaproteobacteria and Alphaproteobacteria (Steps 11,12) [3]. Our results extend the breadth of Form IV RuBisCO distribution to new branches of the tree of life and double the possible instances of its gene transfer, including one across domains. However, it is still unclear which lineage of bacteria among those that are known to bear the IV-Photo enzyme is likely to have been the original source. Similarly, we recovered RuBisCO enzymes related to the Form IV encoded in the genome of at least one Beckwithbacteria and many scaffolds of bacteriophage origin, including one unusually large, manually curated phage genome. One possibility is that a phage acquired a copy of the ancestral Form IV enzyme, termed the DeepYkr [3], from a bacterial ancestor (Step 13), ultimately evolving into a divergent clade. One of these divergent sequences may then have been transferred to Beckwithbacteria (Step 14). The reverse scenario is also possible, in which members of the Myoviridae acquired this enzyme from Beckwithbacteria, possibly as prophages. However, to date, we know of no cases of >200 kbp phage genomes integrating into small (generally < 1 Mbp) CPR genomes. The discovery of RuBisCO homologs in phage also provides a possible mechanism for the widespread lateral gene transfer of this enzyme observable across the tree of life (**Fig. 5**) [28]. However, as of yet, phage encoding ‘bona fide’ RuBisCO forms have not been identified.

The Form II/III enzymes are presently distributed among the CPR, DPANN, and at least one methanogenic archaeal lineage [4,13]. Given that this form is most closely related to Form II (**Fig. 1a**), we speculate here that the Form II/III enzyme evolved in a CPR lineage (possibly the Dojkabacteria) after transfer of a Form II sequence (Step 15). Form II/III could then be transferred to several other CPR lineages and one or more archaeal lineages (Step 16). Notably, no CPR with Form II enzymes have been reported to date.

Finally, there are two possible explanations for the apparent discordance between the phylogenetic pattern showing Forms II and II/III branching together and separate from Form I (**Fig. 1a**) and the pathway association of these Forms (I and II in CBB vs. II/III in the AMP pathway). If Form II/III preserves its ancestral function in the AMP pathway then the CBB pathway function in Forms I and II must have arisen by convergent evolution. Alternatively, the CBB pathway function in Forms I and II shared a common ancestor and Form II/III reverted back to function in the AMP pathway, possibly due to loss of the other CBB pathway enzymes. We suggest that convergent evolution of the more complex CBB pathway (which also requires PRK and transketolase, as well as various glycolysis enzymes) is less likely than reversion due to gene loss, especially given that gene loss is likely to have been widespread in the CPR. Methanogens and DPANN archaea, which generally do not have PRK, could then have acquired the Form II/III RuBisCO by lateral transfer (**Fig. 5**).

### Conclusion

In conclusion, we show that CPR bacteria, DPANN archaea, and bacteriophage harbor RuBisCO diversity that broadens our understanding of the distribution of this enzyme across the tree of life. The wide but sparse distribution of RuBisCO within the CPR and DPANN may be the consequence of extensive lateral gene transfer as well as gene loss. Further, some transfers may have catalyzed major shifts in carbon metabolism in bacterial lineages with limited metabolic repertoires. Specifically, lateral transfer of RuBisCO could have conferred a “missing puzzle piece” for organisms already bearing AMP phosphorylase and R15P isomerase, completing the genomic repertoire for the CO_2_-incorporating AMP metabolism present in extant lineages. Likewise, Gottesmanbacteria may have evolved a partial CBB cycle or augmentation to the AMP pathway by linking laterally acquired RuBisCO and PRK to genes in the oxidative pentose phosphate pathway (**Fig. 4, Fig. S6**). Clearly, metagenomic studies of diverse environments can help to shed light on phylogenetic distribution and also to extend models of evolution for even well-studied enzymes.

## Acknowledgements

We thank Brian Thomas, Shufei Lei, Lin-xing Chen, Adi Lavy, Karthik Anantharaman, Alexander Probst, and Patrick Shih for informatics support and helpful discussions. Funding was provided by the Berkeley Fellowship to A.L.J., the Innovative Genomics Institute at the University of California, Berkeley, and the Chan-Zuckerberg Biohub Initiative. C.L.D. was supported by the Alternative Earths NASA Astrobiology Institute.

## Author contributions

A.L.J. conducted the phylogenetic analyses, A.L.J. and C.J.C performed genomic analyses, J.F.B. carried out genome curation, and C.J.C. conducted the protein modeling. J.F.B., C.J.C., C.L.D. developed the project. A.L.J and J.F.B. wrote the manuscript, and all authors read and made comments on the manuscript prior to submission.

## METHODS

### Genome collection and annotation

We gathered a set of ∼4,000 CPR/DPANN genomes from metagenomes from several previous studies of groundwater, soil, ocean, and subsurface environments. Additionally, we binned new genomes from a sediment from Rifle, CO [29] and the water column in the Baltic Sea [30]. Binning methods and taxonomic assignments followed those described in Anantharaman et al. [29]. Several genome fragments were manually curated, making use of unplaced paired reads to increase their length. This was necessary to test for bacterial affiliation based on comparison of the encoded genes with genes in known CPR genomes.

Proteins were predicted for each genome using Prodigal (“meta” mode) [31]. Preliminary functional predictions were established using a pipeline based on KEGG Orthology [32]. All vs. all global search of proteins in each KO from the KEGG database was performed using usearch [33], and protein percent identity was used as input to MCL clustering [34] with inflation parameter of 1.1. For each resulting cluster, the proteins were aligned using MAFFT version 7 [35], and Hidden Markov Models (HMMs) were constructed using the HMMER suite [36]. Predicted proteins from each CPR/DPANN genome in this study were scanned using hmmsearch [36], and annotation was assigned according to the best HMM hit, providing it was above a pre-defined KEGG Orthology noise cutoff.

### RuBisCO analysis

We extracted above-threshold hits for RuBisCO large chain (K01601), yielding a final set of genomes encoding the enzyme. To analyze the number of non-redundant genomes containing RuBisCO, we repeated the above analysis with a set of ∼3000 high quality genomes from various environments. These genomes were de-replicated at 99% secondary ANI using dRep (-comp 20) [37] and then analyzed for presence of RuBisCO.

To expand the breadth of our main RuBisCO set, we identified RuBisCO sequences (many of which were unbinned) from sediment and groundwater metagenomes [10,29,38]. We excluded sequences shorter than 200 amino acids in length to remove fragmented proteins. Phylum-level taxonomy for these sequences was assigned based on the closest affiliation of the encoded sequences. These sequences were added to those from genomes and the entire set was dereplicated (USEARCH, -id 0.99 -sort length) [33]. Sourcing for de-replicated sequences can be found in **Table S1**. Next, we combined the full set with reference RuBisCO from NCBI and aligned it using MAFFT (default parameters) [35]. Alignments were trimmed by removing columns with > 95% gaps. We constructed a maximum likelihood tree with RaXML (default parameters, cipres.org) and subsequently assigned each RuBisCO sequence to previously identified Forms based on phylogenetic clustering with reference sequences. Binned sequences excluded from the de-replicated set were re-inserted into the tree and classified for downstream analyses. Sequences in ambiguous phylogenetic positions were annotated as “unknown.” Custom HMMs were constructed for each RuBisCO form using the HMMER suite [36] and were subsequently self-tested and manually refined to exclude low-scoring sequences.

### Collection and analysis of viral sequences

To explore the possibility that phage encode RuBisCO, we generated a database of putative phage genome fragments using sequences from IMG/VR (img.jgi.doe.gov/vr/) and several previous metagenomic studies of groundwater and subsurface environments [29,38]. Contigs from the latter metagenomes were assigned a putative phage origin if the majority of encoded genes had no identifiable sequence similarity to genes in bacterial (or archaea). Only sequences >10 kbp in length were included to improve the confidence of phage assignments. Predicted proteins from putative phage contigs were interrogated using the above RuBisCO HMMs, and those with significant HMM hits at or above a score of 100 and e<< 0.05 were retained for further analysis. Genome fragments with RuBisCO-related sequences were further evaluated to confirm the presence of additional (e.g., structural) genes indicative of phage classification. Once manually verified, the putative RuBisCO proteins of phage origin were de-replicated at 99% identity and incorporated into the phylogenetic analysis. Phage terminase proteins were extracted using existing annotations (in the case of IMG/VR fragments) or BLAST-based annotations (in the case of groundwater/subsurface fragments). To establish putative identity, we then aligned recovered phage terminases with reference proteins and created a tree with RaxML. Finally, several additional phage genome fragments encoding RuBisCO-related sequences were manually curated to increase their length.

### Residue analysis and protein modeling

To analyze the biochemically-relevant characteristics of the RuBisCO sequences included in the de-replicated set, including those in the putative phage category, we extracted 12 residues known to be important for catalytic activity and 7 important for substrate binding from each sequence [2,15]. A sequence logo of these 19 sites for each Form was constructed using WebLogo [39]. Additionally, we modeled exemplary Form III, Form III-c, and putative phage proteins using the I-TASSER suite (zhanglab.ccmb.med.umich.edu/I-TASSER/) [40].

### Pathway and contig-level analyses

Finally, we identified two sets of proteins involved in RuBisCO-mediated carbon metabolism (**Table S2**) within the binned genomes using the KEGG annotation results. Of particular interest were AMP phosphorylase (*deoA*) and R15P isomerase (*e2b2*), thought to be involved in AMP metabolism [6], and phosphoribulokinase (PRK), a marker gene for the CBB pathway. We examined the distribution of these genes across genomes as well as their genomic context. Specifically, for the three genes likely involved in AMP metabolism, the proximity of the genes on the same contig was calculated by taking the sum the lengths of the intervals between the genes. Gene fusions were identified by noting abnormally long RuBisCO sequences and examination of the domain structure through NCBI blastp (blast.ncbi.nlm.nih.gov/Blast.cgi). Phylogenetic trees for PRK, *deoA*, and *e2b2* were created using the procedure described above for RuBisCO and visualized using iTOL [41].

### Data and software availability

The newly binned genomes from Rifle Sediment and Landsort Deep (Baltic Sea) are deposited in NCBI under accession number (TBA).

Custom code and intermediate data files used for the described analyses are available in interactive Jupyter Notebook format at https://github.com/alexanderjaffe/cpr_dpann_rubisco/.

## SUPPLEMENTAL INFORMATION

**Figure S1.**
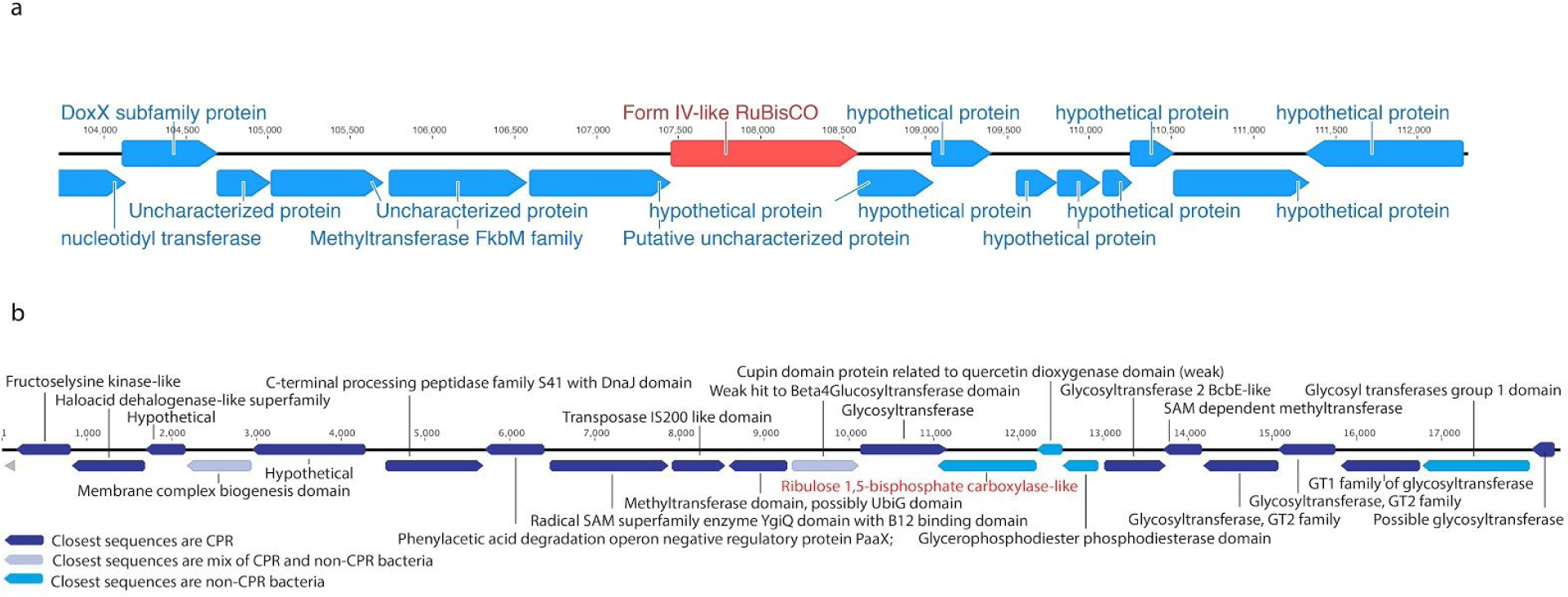
Genomic context of Form IV-like RuBisCO in curated genomes putatively assigned to **(a)** bacteriophage and **(b)** Beckwithbacteria.

**Figure S2.**
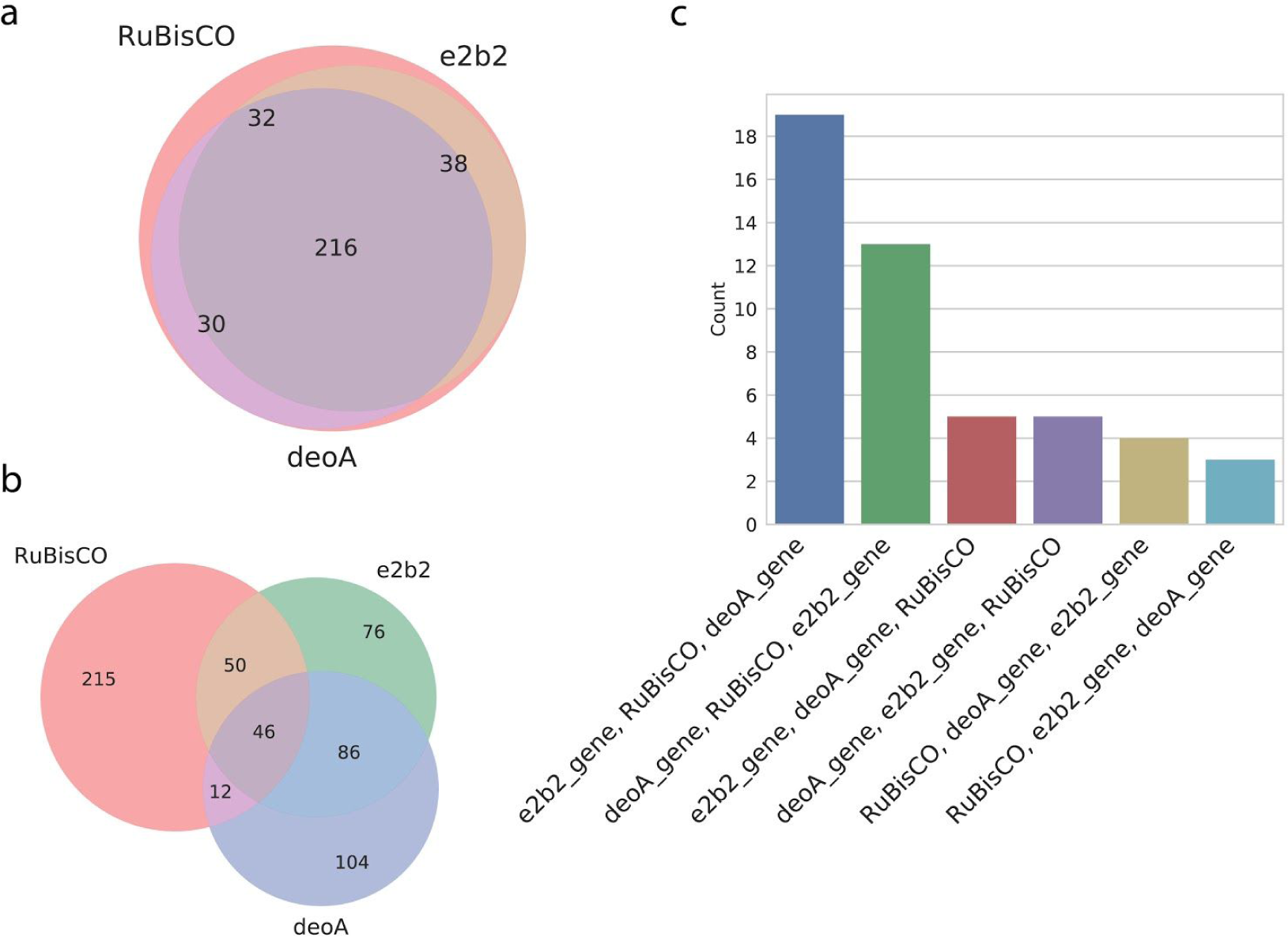
Analysis of AMP pathway components in CPR and DPANN genomes. Venn diagrams describing the number of **(a)** CPR/DPANN genomes and **(b)** assembled genome fragments encoding one, two, or three of the pathway components. **(c)** On genome fragments encoding all three pathway components, the frequency of unique ordering of these components.

**Figure S3.**
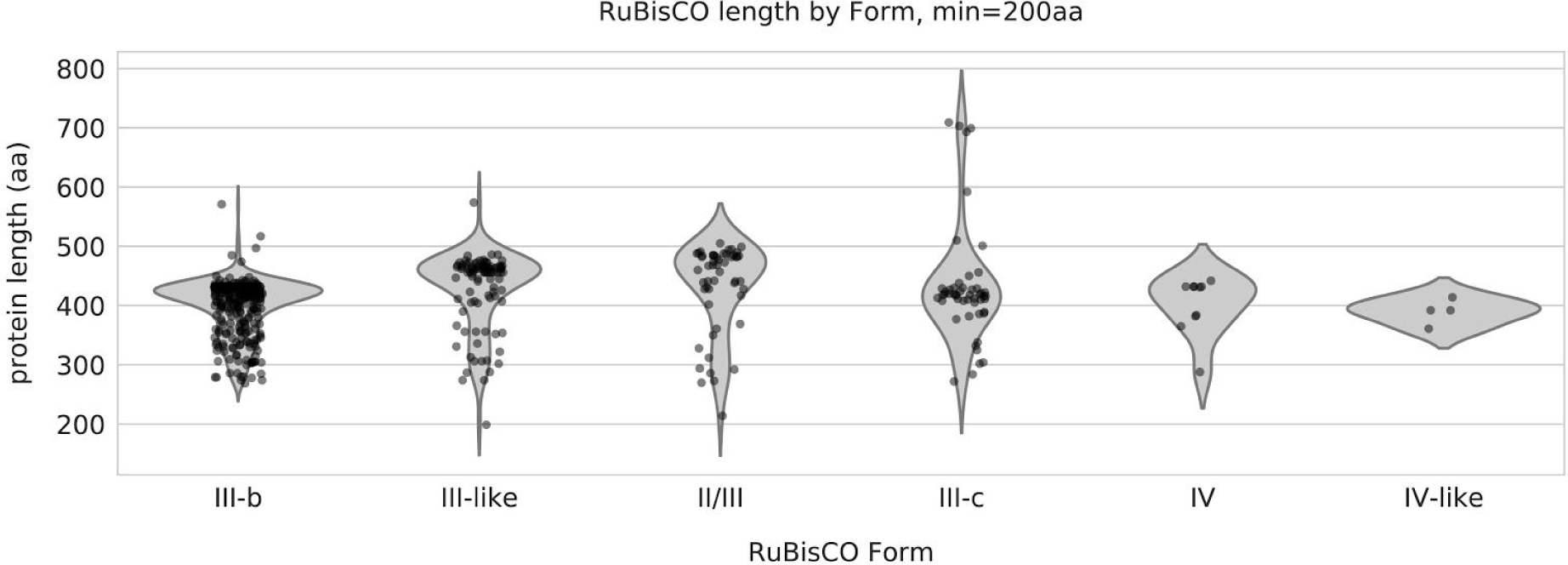
RuBisCO protein length by phylogenetic Form for ∼330 binned CPR/DPANN genomes.

**Figure S4.**
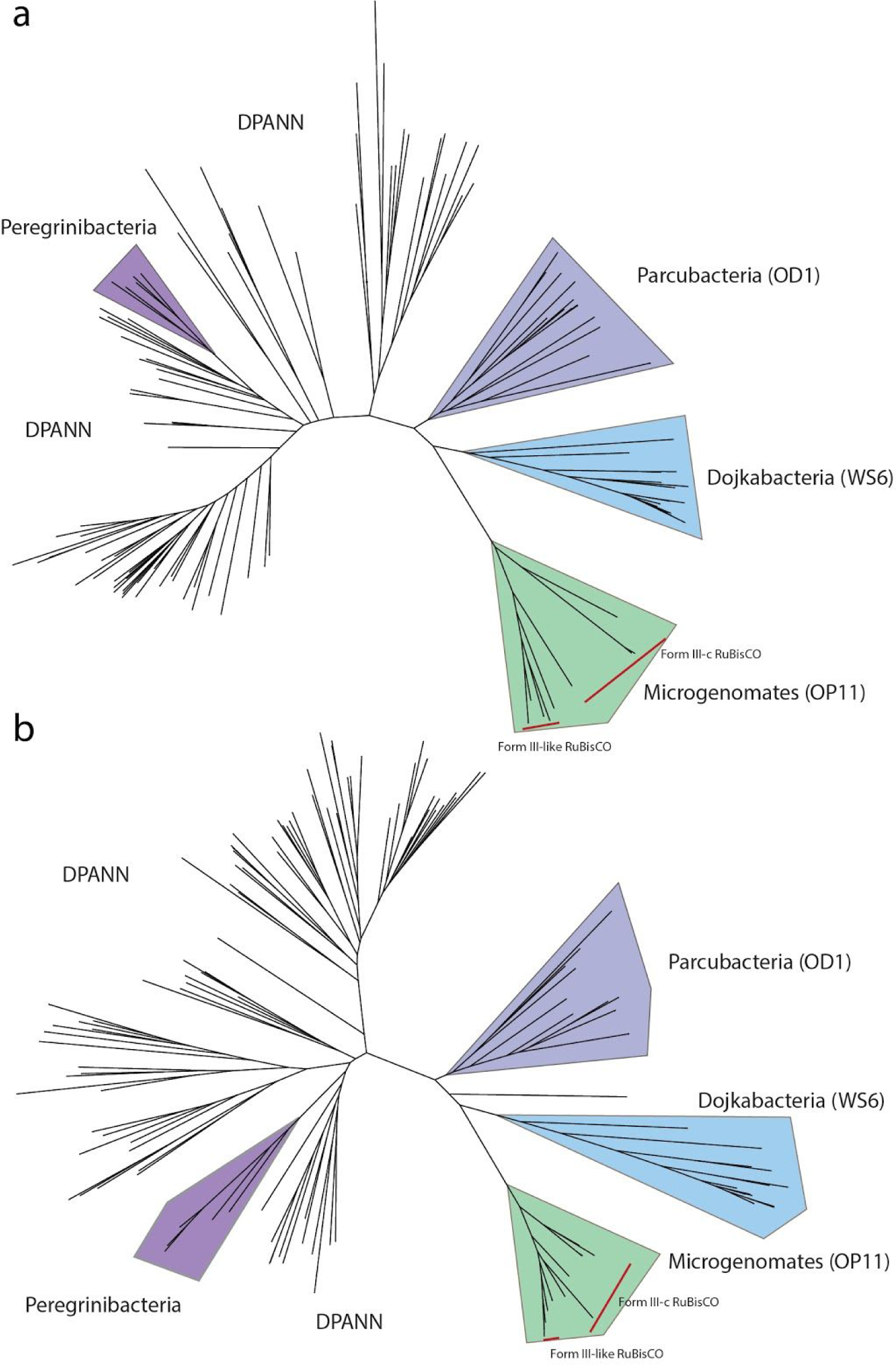
Molecular phylogenies for *e2b2* (R15P isomerase) and *deoA* (AMP phosphorylase) from ∼ 330 CPR/DPANN genomes.

**Figure S5.**
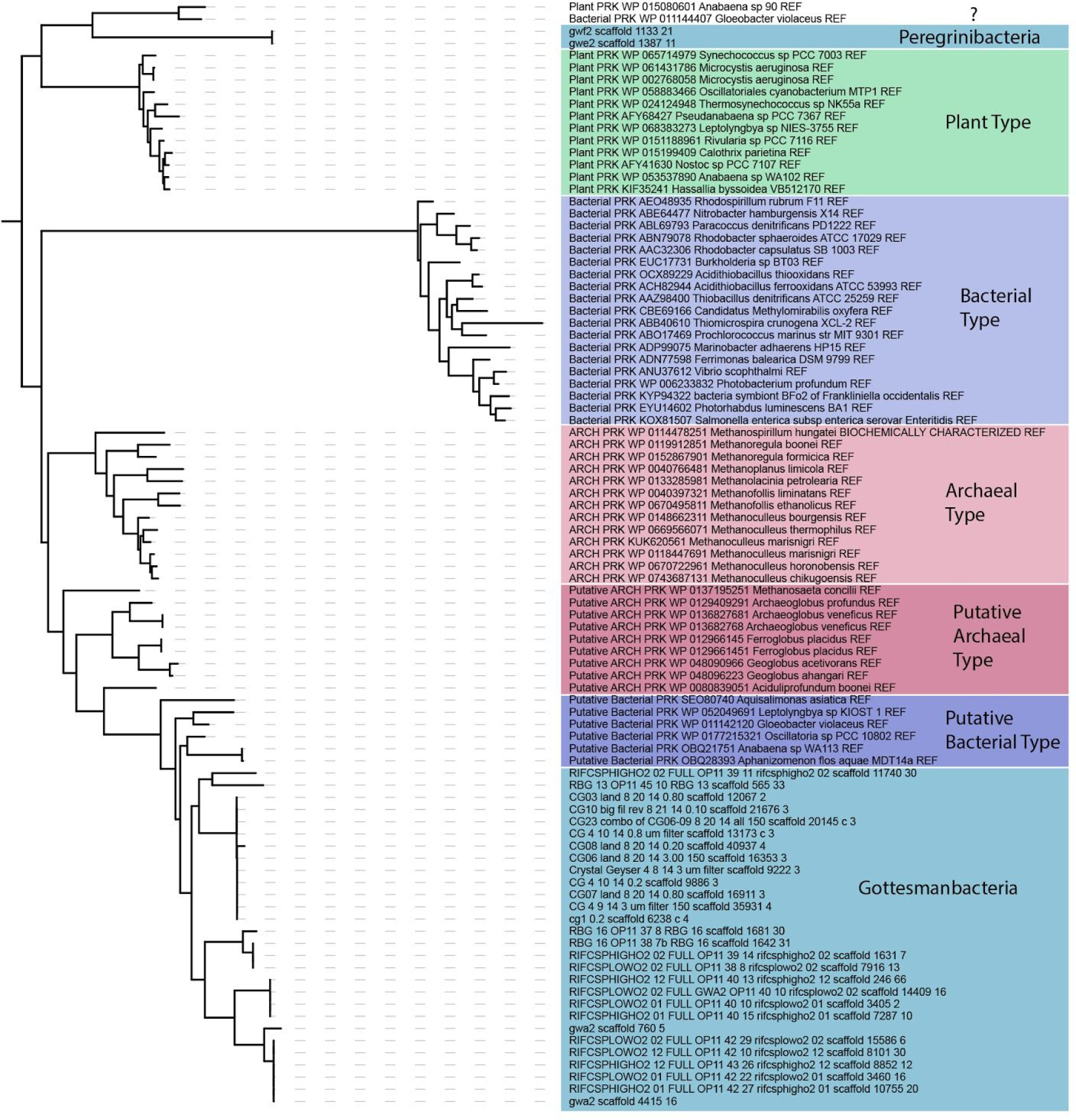
Detailed molecular phylogeny for phosphoribulokinase (PRK) including putative sequences from the CPR phyla Gottesmanbacteria and Peregrinibacteria.

**Figure S6.**
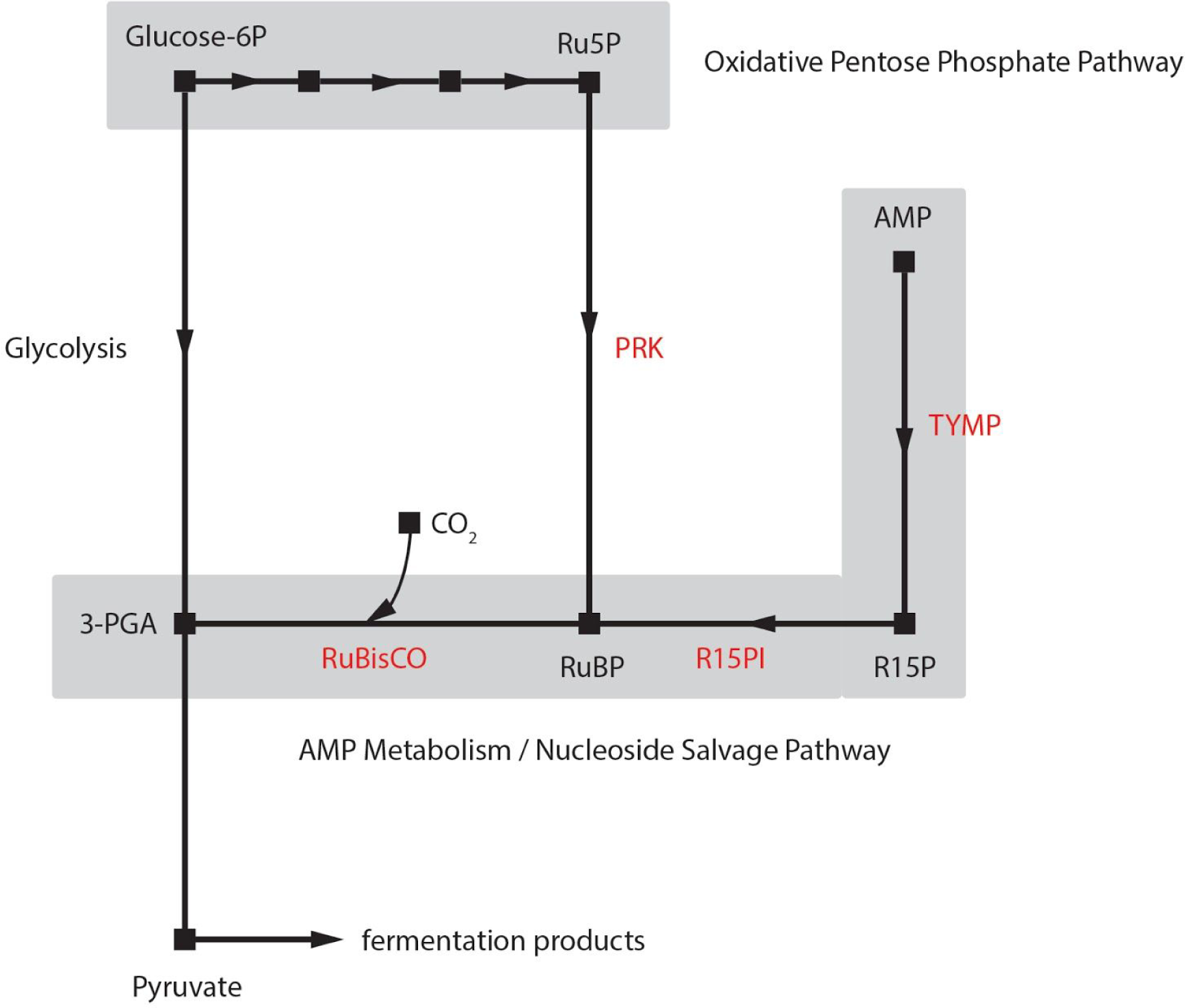
Possible augmentation of carbon metabolism in Gottesmanbacteria by integration of phosphoribulokinase (PRK) with the oxidative pentose phosphate pathway and the AMP pathway. Abbreviations: AMP, adenosine monophosphate; Ru5P, ribulose-5-phosphate; TYMP, AMP phosphorylase; R15P, ribose-1-5-bisphosphate; R15PI, ribose-1,5-bisphosphate isomerase; RuBP, ribulose-1,5-bisphosphate; 3-PGA, 3-phosphoglyceric acid.

**Figure S7.**
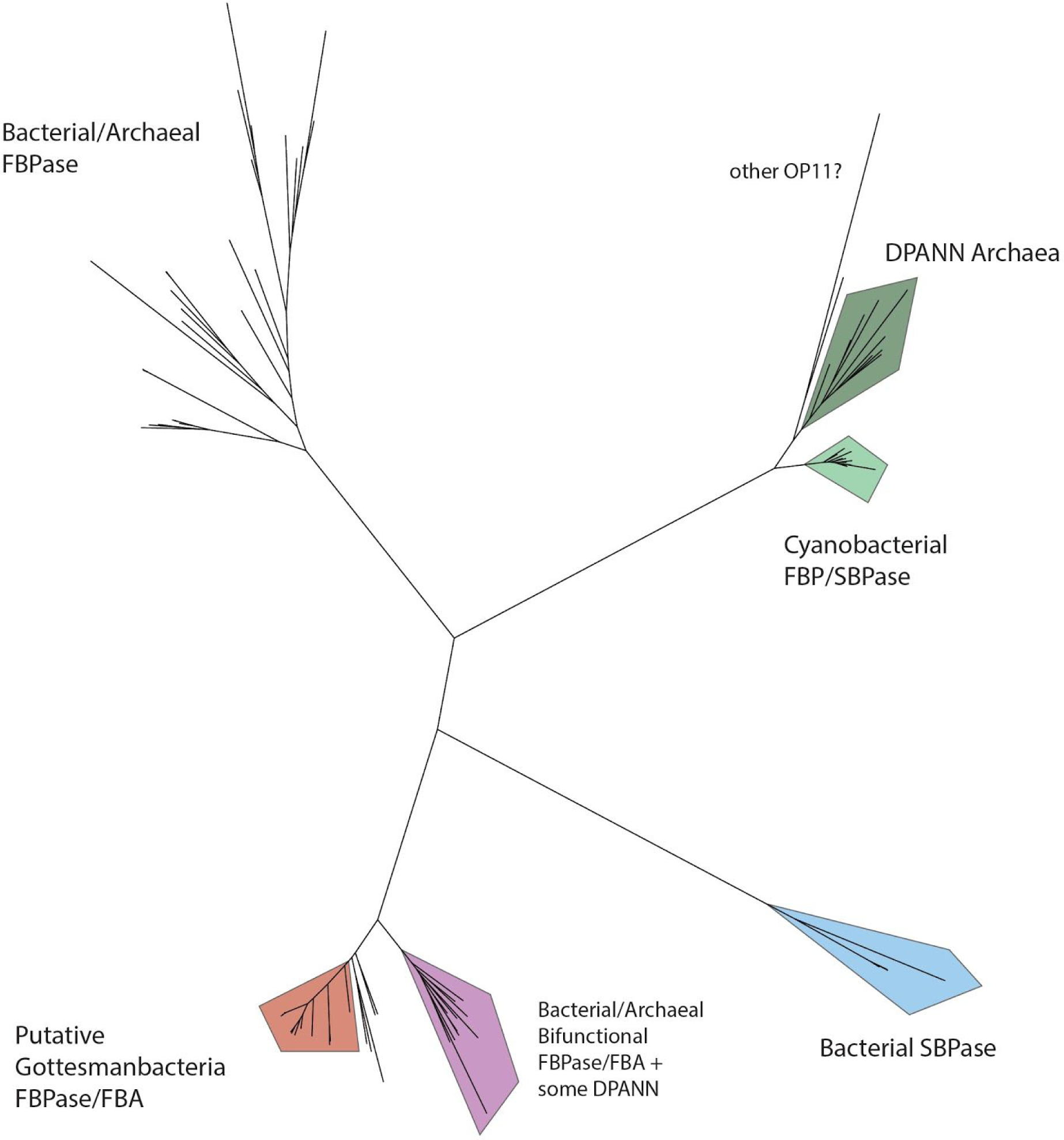
Combined molecular phylogeny for fructose-1,6-bisphosphatase (FBPase), fructose-1,6-bisphosphate aldolase (FBA), sedoheptulose-bisphosphatase (SBPase) for ∼330 CPR and DPANN genomes in addition to various reference forms.

**Table S1.** Sourcing for the de-replicated set of RuBisCO sequences used in phylogenetic analyses.

| Source | # Seqs | Reference(s) |
| --- | --- | --- |
| Rifle Groundwater and Sediment | 375 | Anantharaman et al., 2016 (doi:10.1038/ncomms13219) |
| Crystal Geyser (Groundwater) | 105 | Probst et al., 2017 (doi:10.1038/s41564-017-0098-y), Probst et al., 2017 (doi:10.1111/1462-2920.13362 ) |
| IMG/VR | 79 | img.jgi.doe.gov/vr/ |
| Other | 36 | n/a |
| "Uncultivated Bacteria and Archaea" | 22 | Parks et al., 2017 (doi:10.1038/s41564-017-0012-7) |
| Borehole Japan (Subsurface) | 16 | Hernsdorf et al., 2017 (doi:10.1111/1462-2920.13362 ) |
| TARA Oceans | 10 | Tully et al., 2018 (doi:10.1038/s41564-017-0012-7) |
| NCBI Genome | 5 | ncbi.nlm.nih.gov/genome |
| IMG/MG | 2 | img.jgi.doe.gov |

**Table S2.** KEGG Orthology numbers (KOs) for enzymes involved in the AMP pathway and the Calvin-Benson-Bassham Cycle. See https://www.genome.jp/kegg/.

| Pathway | Enzyme Name | KO(s) |
| --- | --- | --- |
| **AMP Metabolism**<br>**(nucleotide salvage)** | Ribulose-bisphosphate carboxylase (RuBisCO) long chain | K01601 |
|  | AMP phosphorylase (deoA) | K18931, K00758 |
|  | R15P Isomerase (e2b2) | K18237, K08963, K03680 |
| **Calvin-Benson-Bassham Cycle** | Ribulose-bisphosphate carboxylase (RuBisCO) long chain | K01601 |
|  | Phosphoglycerate kinase (PGK) | K00927 |
|  | Glyceraldehyde 3-phosphate dehydrogenase (GAP-DH) | K00134, K05298, K00150 |
|  | Triosephosphate isomerase (TIM) | K01803 |
|  | Fructose-bisphosphate aldolase (FBA) | K01622, K01623, K01624, K11645, K16305, K16306 |
|  | Fructose-bisphosphatase (FBPase) | K01086, K01622, K02446, K03841, K04041, K11532 |
|  | Transketolase (TKL) | K00615 |
|  | Sedoheptulose-bisphosphatase (SBPase) | K01086, K01100, K11532 |
|  | Ribose-5-phosphate isomerase (RPI) | K01807, K01808 |
|  | Ribulose-phosphate 3-epimerase (RPE) | K01783 |
|  | Phosphoribulokinase (PRK) | K00855 |

